# Evolution of an enzyme conformational ensemble guides design of an efficient biocatalyst

**DOI:** 10.1101/2020.03.19.999235

**Authors:** Aron Broom, Rojo V. Rakotoharisoa, Michael C. Thompson, Niayesh Zarifi, Erin Nguyen, Nurzhan Mukhametzhanov, Lin Liu, James S. Fraser, Roberto A. Chica

## Abstract

The creation of artificial enzymes is a key objective of computational protein design. Although *de novo* enzymes have been successfully designed, these exhibit low catalytic efficiencies, requiring directed evolution to improve activity. Here, we used room-temperature X-ray crystallography to study changes in the conformational ensemble during evolution of the designed Kemp eliminase HG3 (*k*_cat_/*K*_M_ 160 M^−1^s^−1^). We observed that catalytic residues were increasingly rigidified, the active site became better pre-organized, and its entrance was widened. Based on these observations, we engineered HG4, an efficient biocatalyst (*k*_cat_/*K*_M_ 120,000 M^−1^s^−1^) containing active-site mutations found during evolution but not distal ones. HG4 structures revealed that its active site was pre-organized and rigidified for efficient catalysis. Our results show how directed evolution circumvents challenges inherent to enzyme design by shifting conformational ensembles to favor catalytically-productive sub-states, and suggest improvements to the design methodology that incorporate ensemble modeling of crystallographic data.

## Introduction

Enzymes are the most efficient catalysts known, accelerating chemical reactions by up to 26 orders of magnitude^1^ while displaying unmatched selectivity. The ability to create, from scratch, an efficient artificial enzyme for any desired chemical reaction (i.e., a *de novo* enzyme) is a key objective of computational protein design. Progress towards this goal has been made over the past few decades following the development of computational enzyme design algorithms.^2,3^ These methods have been used to create *de novo* enzymes for a variety of model organic transformations including the Kemp elimination,^4,5^ retro-aldol,^6,7^ Diels-Alder,^8^ ester hydrolysis,^9^ and Morita-Baylis-Hilman^10^ reactions. Although successful, catalytic activities of *de novo* enzymes have been modest, with *k*_cat_/*K*_M_ values being several orders of magnitude lower than those of natural enzymes.^11,12^ In addition, structural analyses of designed enzymes have revealed important deficiencies in the computational methodologies, resulting in inaccurate predictions of catalytic and ligand-binding interactions,^5^ and thereby low success rates,^4,6,8^ emphasizing the need for continued development of robust enzyme design algorithms.

To improve the catalytic activity of designed enzymes, researchers have used directed evolution. This process has yielded artificial enzymes displaying catalytic efficiencies approaching those of their natural counterparts, and provided valuable information about the structural determinants of efficient catalysis.^4,13–15^ During evolution, active-site residues, including designed catalytic amino acids, were often mutated, leading to enhanced catalysis via optimization of catalytic contacts, ligand binding modes, and transition-state complementarity.^13–15^ Directed evolution has also yielded beneficial mutations at positions remote from the active site. Distal mutations have been shown to enhance catalysis by shifting the populations of conformational sub-states that enzymes sample on their energy landscape towards those that are more catalytically active.^16–18^ Therefore, a better understanding of enzyme conformational ensembles, including the effect of mutations on the population of sub-states, could provide valuable insights to aid in the development of robust computational enzyme design methodologies.

Here, we study changes in the conformational ensemble along the evolutionary trajectory of the *de novo* Kemp eliminase HG3 (*k*_cat_/*K*_M_ 160 M^−1^s^−1^) using room-temperature X-ray crystallography. We observe that during evolution, catalytic residues were increasingly rigidified through improved packing, the active site became better pre-organized to favor productive binding of the substrate, and the active-site entrance was widened to facilitate substrate entry and product release. Based on these observations, we generated a variant that contains all mutations necessary to establish these structural features, which are found at positions within or close to the active site. This variant, HG4, is >700-fold more active than HG3, with a catalytic efficiency on par with that of the average natural enzyme (*k*_cat_/*K*_M_ 120,000 M^−1^s^−1^). Crystallographic analysis of HG4 reveals that mutations proximal to the active site are sufficient to alter the conformational ensemble for enrichment of catalytically-competent sub-states. Lastly, we demonstrate that HG4 can be successfully designed using a crystallographically-derived ensemble of backbone templates approximating conformational flexibility, but not with the single template used to design HG3, offering insights for improving enzyme design methodologies.

## Results

### HG series of Kemp eliminases

Perhaps the most successful example of the improvement of a *de novo* enzyme by directed evolution has been the engineering of HG3.17,^15^ the most active Kemp eliminase reported to date.^11^ This artificial enzyme, which catalyzes the deprotonation of 5-nitrobenzisoxazole into the corresponding *o*-cyanophenolate (Figure 1a), was evolved from the *in silico* design HG3^5^ over 17 rounds of mutagenesis and screening that also yielded the HG3.3b, HG3.7, and HG3.14 intermediates (Figure 1b, Supplementary Table 1). A total of 17 mutations were introduced into HG3 during evolution to produce HG3.17, resulting in a >1000-fold increase in catalytic efficiency (Table 1, Supplementary Figure 1). Of these mutations, 11 occurred at positions within or close to the active site, including 8 at positions that were optimized during computational design of HG3 (Table 1). One of the key active-site mutations occurred at position 50, which was mutated twice during evolution, first from lysine to histidine (HG3 to HG3.3b) and then from histidine to glutamine (HG3.3b to HG3.7), resulting in a novel catalytic residue ideally positioned for stabilizing negative charge buildup on the phenolic oxygen at the transition state (Figure 1a). Comparison of the crystal structure of the earlier *in silico* design HG2 (Supplementary Figure 2) with that of a double mutant of HG3.17, in which surface mutations N47E and D300N were reverted to the corresponding amino acids found in HG3 to facilitate crystallization (HG3.17-E47N/N300D, PDB ID: 4BS0),^15^ revealed that catalytic activity was also enhanced via optimized alignment of the transition-state analogue (TSA) with the catalytic base Asp127 (Figure 1c), and improved active-site complementarity to this ligand (Figure 1d). Given that subtle changes to the conformational ensemble of an enzyme can lead to significant rate enhancements,^16–18^ it is possible that mutations in HG3.17 also contributed to enhanced catalytic efficiency by altering the conformational landscape to enrich catalytically-competent sub-states. However, the structures of HG2 and HG3.17-E47N/N300D were solved in the presence of bound TSA and at cryogenic temperatures, which could have shifted the conformational ensemble towards a single predominant sub-state, thereby limiting our ability to evaluate changes to the conformational landscape during directed evolution.

**Figure 1.**
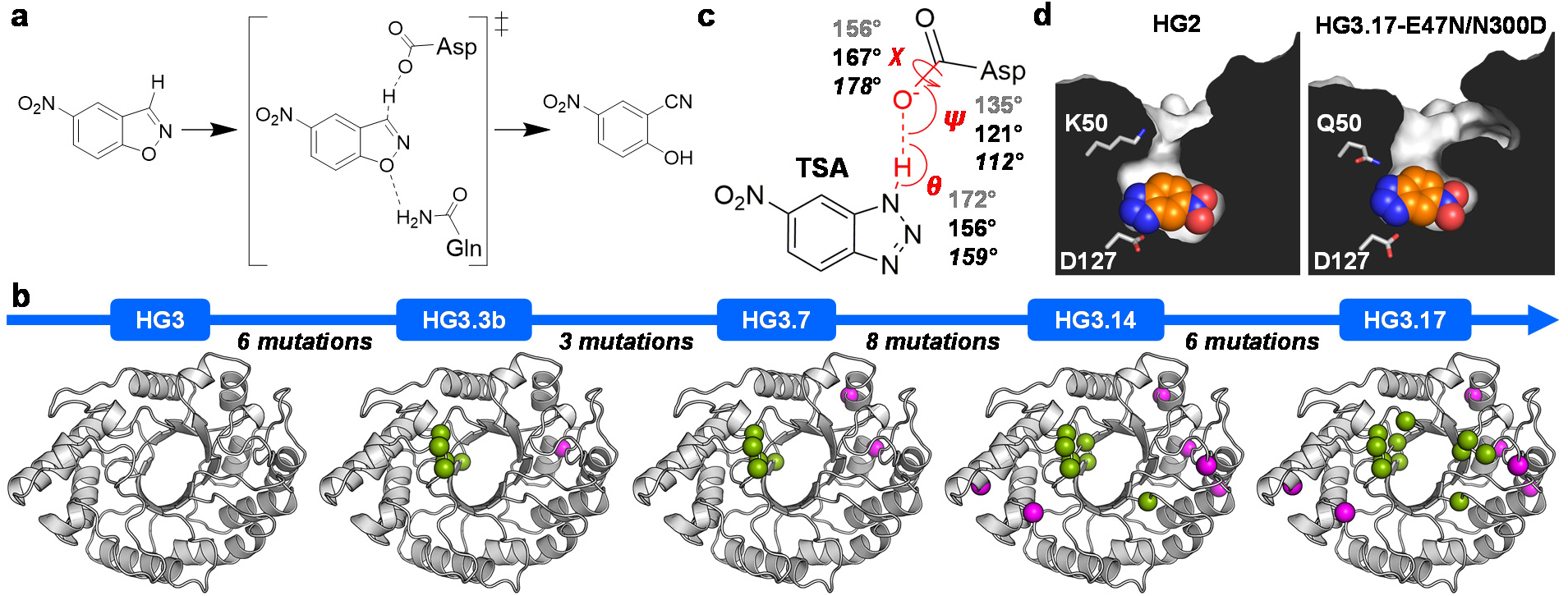
HG series of Kemp eliminases. (a) HG enzymes catalyze the Kemp elimination reaction using a catalytic dyad consisting of a base (Asp127) that deprotonates 5-nitrobenzisoxazole, and an H-bond donor (Gln50) that stabilizes negative charge buildup on the phenolic oxygen at the transition state (‡). This reaction yields 4-nitro-2-cyanophenol. (b) Directed evolution of the *in silico* design HG3, which itself is a single mutant (S265T) of the earlier design HG2. A total of 17 mutations (shown as spheres) were introduced during evolution, including 11 at positions within or close to the active site (green) and 6 at distal sites (magenta). (c) Angles describing the hydrogen bonding interaction between the transition state analog (TSA) and Asp127 in the HG2 (PDB ID: 3NYD)^5^ and HG3.17-E47N/N300D (PDB ID: 4BS0)^15^ crystal structures are indicated in grey and black, respectively. Values in italics are optimal angles calculated for hydrogen bonding interactions between acetamide dimers.^44^ (d) Cut-away view of the active site pocket shows that its structural complementarity with the TSA (spheres) is improved in the higher activity variant HG3.17-E47N/N300D. Key active site residues are shown as sticks.

**Table 1.**
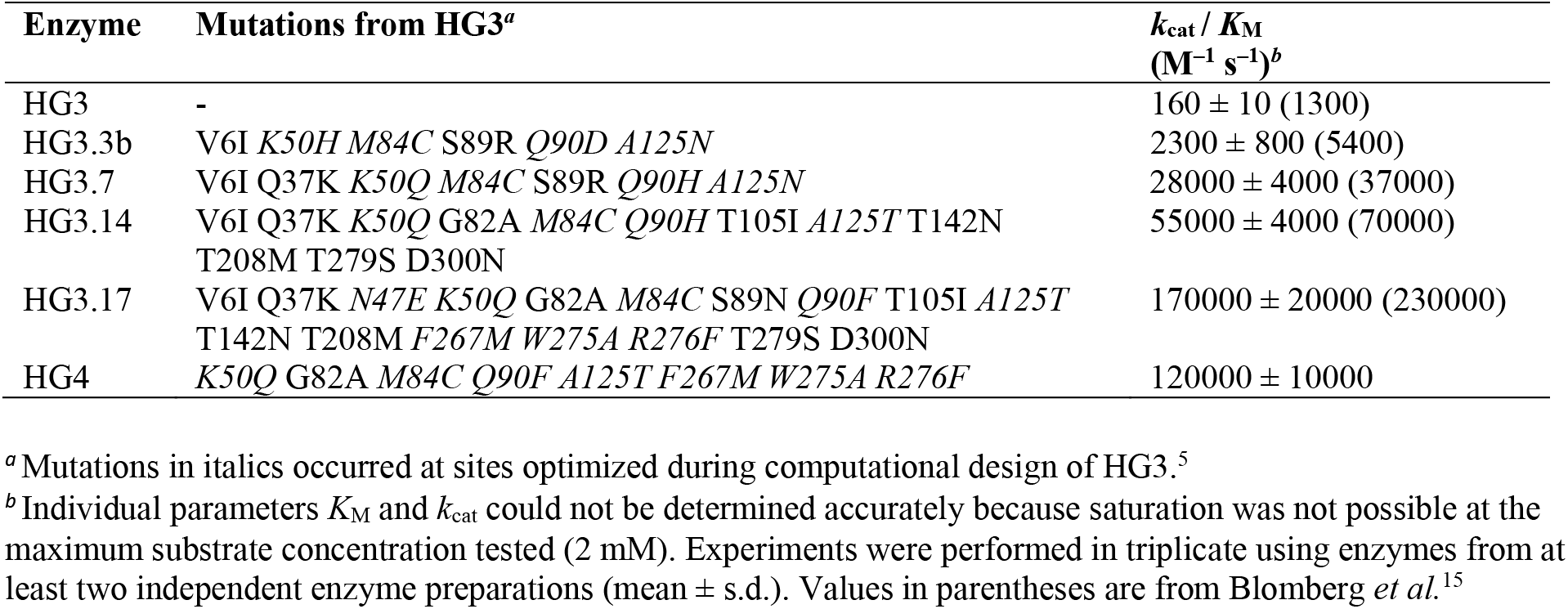
Kinetic parameters of Kemp eliminases.

| Enzyme | Mutations from HG3<sup>a</sup> | $k_{\text{cat}} / K_{\text{M}}$<br>($\text{M}^{-1} \text{s}^{-1}$)<sup>b</sup> |
| --- | --- | --- |
| HG3 | - | 160 ± 10 (1300) |
| HG3.3b | V6I *K50H M84C* S89R *Q90D A125N* | 2300 ± 800 (5400) |
| HG3.7 | V6I Q37K *K50Q M84C* S89R *Q90H A125N* | 28000 ± 4000 (37000) |
| HG3.14 | V6I Q37K *K50Q* G82A *M84C Q90H* T105I *A125T* T142N<br>T208M T279S D300N | 55000 ± 4000 (70000) |
| HG3.17 | V6I Q37K *N47E K50Q* G82A *M84C* S89N *Q90F* T105I *A125T*<br>T142N T208M *F267M W275A R276F* T279S D300N | 170000 ± 20000 (230000) |
| HG4 | *K50Q* G82A *M84C Q90F A125T F267M W275A R276F* | 120000 ± 10000 |

<sup>a</sup> Mutations in italics occurred at sites optimized during computational design of HG3.<sup>5</sup>
<sup>b</sup> Individual parameters $K_{\text{M}}$ and $k_{\text{cat}}$ could not be determined accurately because saturation was not possible at the maximum substrate concentration tested (2 mM). Experiments were performed in triplicate using enzymes from at least two independent enzyme preparations (mean ± s.d.). Values in parentheses are from Blomberg *et al.*<sup>15</sup>

### Room-temperature crystal structures

To evaluate changes to the HG3 conformational ensemble along its evolutionary trajectory, we solved room-temperature (277 K) X-ray crystal structures of all HG-series Kemp eliminases, both in the presence and absence of bound TSA. Room-temperature X-ray crystallography can reveal conformational heterogeneity in protein structures that would not be visible at cryogenic temperatures and thereby provide insights into the conformational ensemble that is sampled by a protein in solution.^19^ All five enzymes yielded crystals under similar conditions (Supplementary Table 2), and these diffracted at resolutions of 1.35–1.99 Å (Supplementary Table 3). All unit cells corresponded to space group P2_1_2_1_2_1_ with two protein molecules in the asymmetric unit, except that of HG3.17, whose asymmetric unit was half the volume of the others and contained only one polypeptide chain although the space group was also P2_1_2_1_2_1_. This result is in contrast with the deposited structure of HG3.17-E47N/N300D, which contains two molecules in a unit cell of identical space group and similar dimensions to those of all other HG variants reported here.^15^ This discrepancy between our structure of HG3.17 and the previously published structure of HG3.17-E47N/N300D is likely caused by the presence of the Asn47 surface residue in all variants except for HG3.17, since this amino acid is involved in crystal packing interactions.

All HG-series enzymes bound the TSA in the same catalytically productive pose (Figure 2a) as that observed in HG2 and HG3.17-E47N/N300D (Figure 1c–d). In this pose, the acidic N– H bond of the TSA that mimics the cleavable C–H bond of the substrate is located within hydrogen-bonding distance to the carboxylate oxygen of Asp127 (2.5–2.6 Å distance between heavy atoms), while the basic nitrogen atom corresponding to the phenolic oxygen of the transition state forms an H-bond with either a water molecule (HG3), the Nε atom of His50 (HG3.3b), or the side-chain amide nitrogen of Gln50 (HG3.7, HG3.14, HG3.17). In addition to being held in place by these polar interactions, the TSA is sandwiched between the hydrophobic side chains of Trp44 and Met237 (Figure 2b), which are part of a binding pocket that also includes the side chains of Ala21, Met/Cys84, Met172, Leu236, Thr265, and Phe/Met267, as well as the backbone of Gly83 and Pro45 (Supplementary Figure 3). Interestingly, the *cis* peptide bond formed between residues 83 and 84 that is present in the original *Thermoascus aurantiacus* xylanase 10A template used to design HG3 (PDB ID: 1GOR^20^) is maintained in all HG structures (Figure 2c), even though both residues were mutated to obtain HG3 (H83G and T84M). In addition to adopting a *cis* conformation, which is stabilized by hydrogen bonding to an ordered water molecule, this peptide bond also adopts the *trans* conformation in the structures of TSA-bound HG3 and HG3.3b. However, starting at HG3.7, the peptide bond is found exclusively in the *cis* conformation in the TSA-bound structures because it is stabilized by an additional hydrogen bond with the Gln50 side-chain carbonyl oxygen. This hydrogen bonding interaction helps to lock Gln50 in a conformation that is properly oriented to stabilize negative charge buildup on the phenolic oxygen at the transition state, likely accounting for the majority of the 12-fold catalytic efficiency enhancement observed in HG3.7 relative to HG3.3b (Table 1).

**Figure 2.**
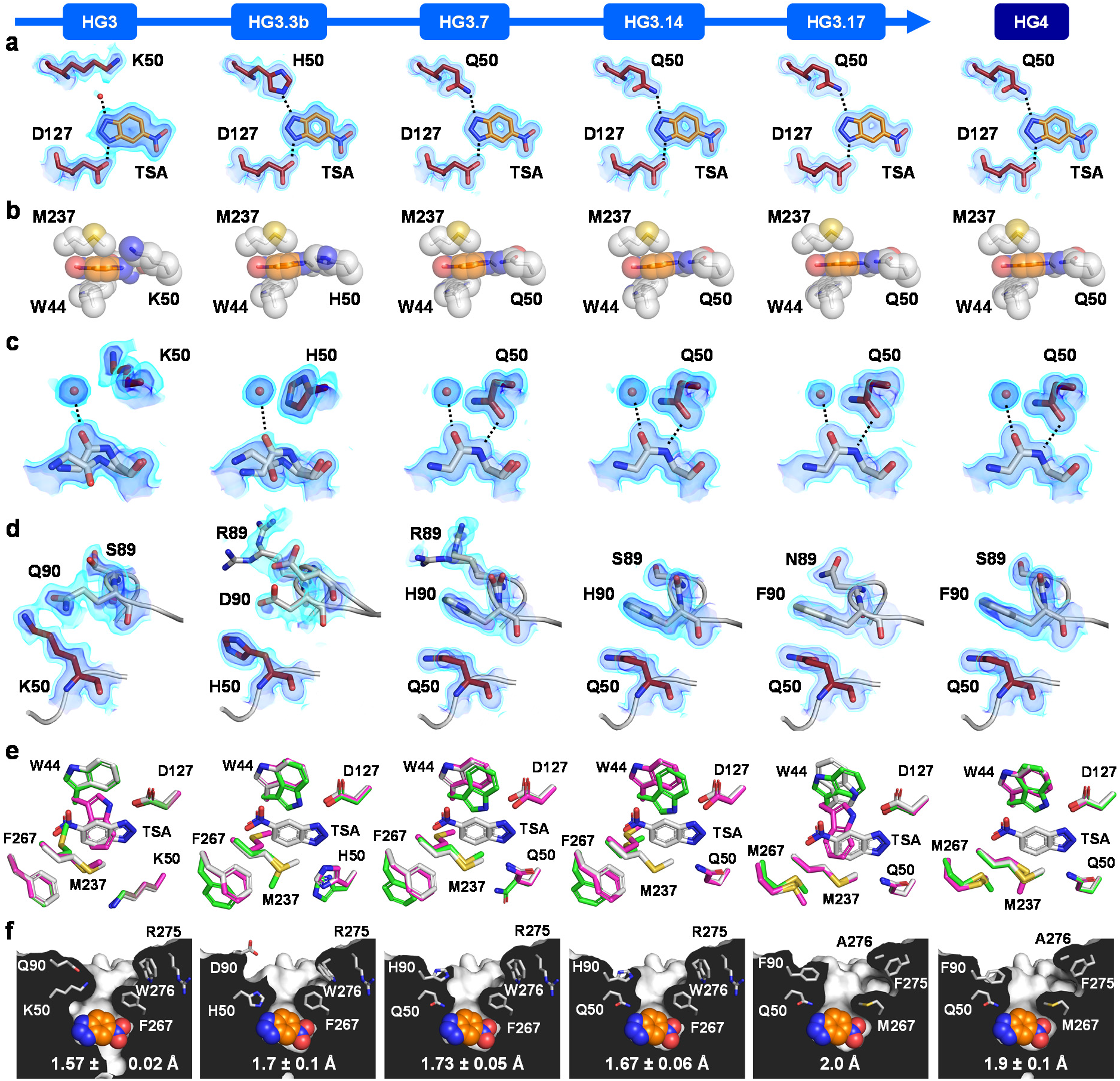
Crystal structures of HG-series Kemp eliminases. In all cases, only atoms from chain A are shown. (a) Binding pose of the transition state analogue (TSA, orange). Hydrogen bonds are shown as dashed lines. The red sphere represents a water molecule. The 2Fo-Fc map is shown in volume representation at two contour levels: 0.5 eÅ^−3^ and 1.5 eÅ^−3^ in light and dark blue, respectively. (b) The TSA (orange) is sandwiched between the hydrophobic side chains of Trp44 and Met237. (c) The peptide bond between residues 83 and 84 can adopt *cis* or *trans* conformations. Hydrogen bonds are shown as dashed lines. The 2Fo-Fc map is shown in volume representation at two contour levels: 0.5 eÅ^−3^ and 1.5 eÅ^−3^ in light and dark blue, respectively. (d) Conformational changes to loop formed by residues 87– 90 over the course of the evolutionary trajectory. The 2Fo-Fc map is shown in volume representation at two contour levels: 0.50 eÅ^−3^ and 1.5 eÅ^−3^ in light and dark blue, respectively. (e) Superposition of the TSA-bound structure (white) with the highest (magenta) and lowest (green) occupancy conformers of the unbound structure for each Kemp eliminase. The occupancies of non-productive conformers of Trp44 in the unbound structures of HG3.17 and HG4 are 62% and 26%, respectively. (f) Cut-away view of the active site shows that its entrance (top) becomes widened during evolution, as indicated by an increasing bottleneck radius (reported as the average radius ± s.d. calculated using the highest occupancy conformers from both chain A and B, except for HG3.17, which contains a single chain). The TSA is shown as orange spheres. Bottleneck radii were calculated using the PyMOL plugin Caver 3.0.^21^

From HG3.7 to HG3.17, no further changes in catalytic residues occurred during evolution. Yet, the catalytic efficiency increased by approximately 6-fold (Table 1). To evaluate whether this increase in activity was caused by changes to the conformational ensemble, we analyzed the B-factors of catalytic residues, which can be interpreted as a measure of the average displacement of an atom, or group of atoms, in the crystal. Since both conformational heterogeneity and crystalline disorder can contribute to atomic B-factors, with the latter effect potentially varying between different crystals, we calculated the Z-scores of the atomic B-factors and compared those across our crystal structures of different HG variants. This Z-score analysis allowed us to evaluate the variation of B-factors relative to the mean value within an individual crystal, and showed that rigidity of the Asp127 side chain did not vary significantly during evolution (Figure 3a). By contrast, the side chain of residue 50 became increasingly rigidified over the course of the evolutionary trajectory. Increasing rigidity at position 50 is expected when this residue is mutated from a lysine to a histidine (HG3 to HG3.3b), given the lower number of degrees of freedom in the latter amino acid. This trend is also expected when histidine at position 50 is mutated to a glutamine (HG3.3b to HG3.7) given the ability of glutamine but not histidine to hydrogen-bond with the *cis* peptide formed by residues Gly83 and Cys84 (Figure 2c). However, rigidity continues to increase at this position between HG3.7 and HG3.17, even though the side-chain rotamer of Gln50 in the presence of bound TSA remains the same (Figure 2a). This result suggests that other structural features contribute to the increased rigidity observed at this position.

**Figure 3.**
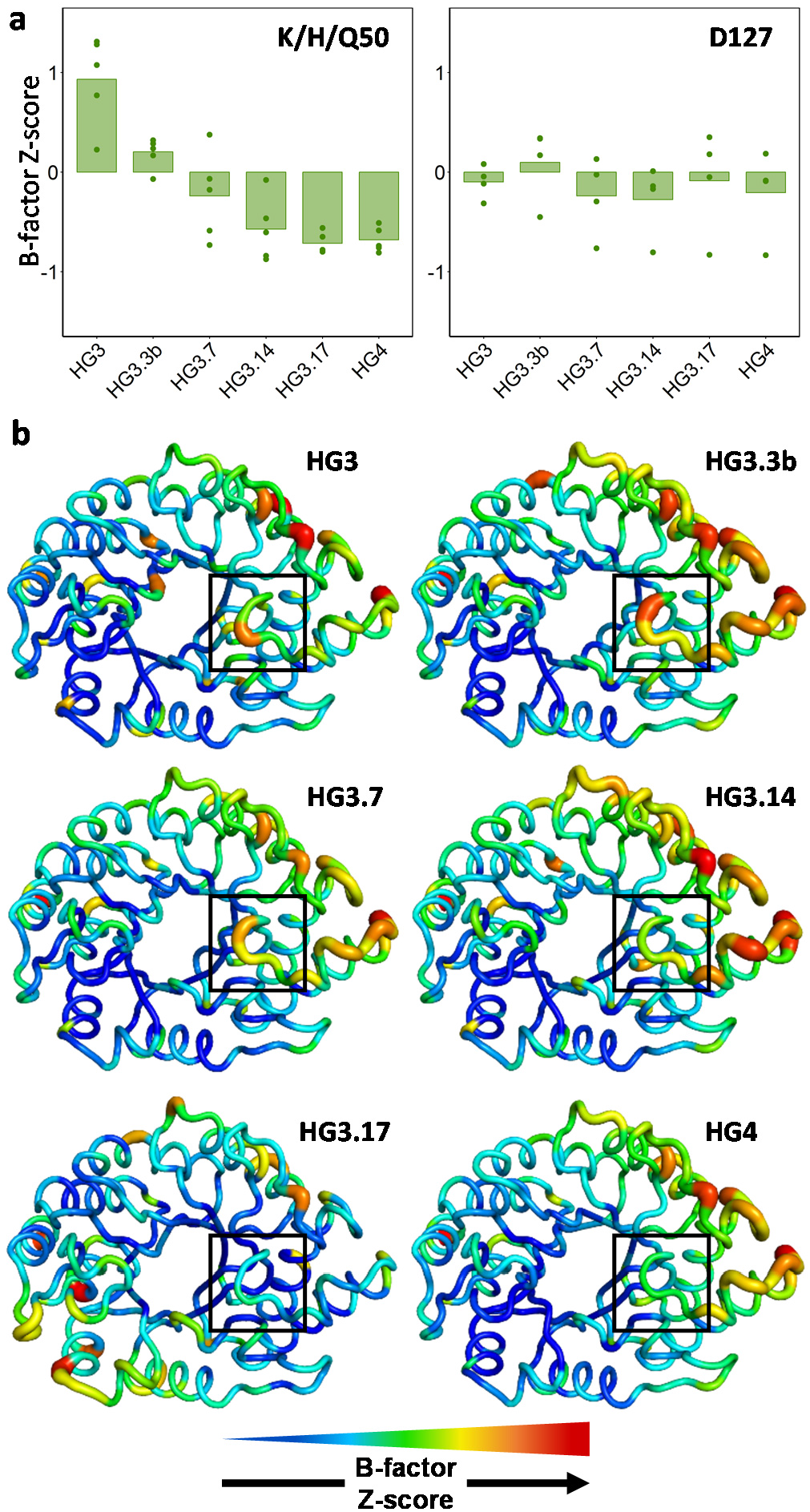
Conformational heterogeneity. (a) B-factor Z-scores for the residue at position 50 in the absence of bound TSA decrease over the course of the evolutionary trajectory, while those for Asp127 do not change significantly. Z-scores of individual side-chain atoms are shown as dots, while the average is indicated by the bar. Positive and negative Z-scores indicate increased flexibility or rigidity relative to the average residue in the protein, respectively. (b) B-factor Z-scores for all protein residues in the absence of bound TSA plotted on a model backbone for each Kemp eliminase. Thickness of the sausage plot increases with the B-factor Z-score, indicating increased flexibility. The loop formed by residues 87–90 (boxed) becomes more rigid during evolution.

To verify the underlying cause of the increased rigidity at position 50, we calculated the average Z-score of atomic B-factors for each residue. We observed a trend whereby the loop formed by residues 87–90, which is located directly on top of residue 50, becomes increasingly rigidified during evolution (Figure 3b). Interestingly, two residues forming this loop (89 and 90) were mutated multiple times over the course of the evolutionary trajectory (Table 1). These mutations induce a conformational change in the loop that moves it closer to the active site, which results in a pi-stacking interaction between the phenyl and carboxamide groups of Phe90 and Gln50 that increases rigidity of the catalytic residue (Figure 2d, Supplementary Figure 4a).

A key determinant of efficient enzyme catalysis is active site pre-organization, which enables enzymes to bind substrates in a geometry close to that of the transition state. To evaluate changes in active site pre-organization during evolution, we compared the structures of HG-series Kemp eliminases in the presence and absence of bound TSA. In all enzymes except for HG3.17, the unbound state is never pre-organized for catalysis as both Trp44 and Met237 adopt conformations that would prevent productive binding of the TSA (Figure 2e). In addition, the His50 and Gln50 catalytic residues in HG3.3b and HG3.7, respectively, adopt a low-occupancy, catalytically non-productive conformation in the unbound state that cannot interact favorably with the TSA. Interestingly, the non-productive conformation of Gln50 in the HG3.7 unbound state (26% occupancy) cannot stabilize the *cis* peptide bond formed by residues 83 and 84 via a hydrogen bonding interaction, and accordingly, the *trans* peptide conformation is also observed in this structure (25% occupancy) (Supplementary Figure 4b).

In contrast with all other HG variants, the unbound state of HG3.17 is correctly pre-organized for catalysis in a large portion of the molecules in the crystal, with only Trp44 adopting a non-productive conformation at 62% occupancy (Figure 2e). In this variant, Met237 adopts exclusively the productive conformer in the unbound state, which is stabilized by packing interactions with the neighboring Met267 side chain, a mutation that was introduced late in the evolutionary trajectory (HG3.14 to HG3.17). Overall, three of the four residues that are key for binding and stabilizing the TSA (Gln50, Asp127, Met237) adopt a catalytically productive conformation in the HG3.17 unbound state, resulting in approximately 40% of the molecules in the crystal being correctly pre-organized for efficient catalysis.

Enhanced complementarity to the transition state is another important feature of efficient catalysis. Therefore, computational enzyme design algorithms aim to optimize packing of the transition state. However, transition-state overpacking may reduce catalytic efficiency by creating a high-energy barrier preventing substrate entry and product release. To evaluate whether active-site accessibility changed during evolution, we calculated the active-site entrance bottleneck radius on TSA-bound structures.^21^ We observed that during evolution, the active-site bottleneck formed by the side chains of residues 50 and 267, became widened (Figure 2f), as did the mouth of the substrate entry channel formed by residues Arg275 and Trp276, which were mutated to smaller amino acids. This widening of the active site entrance could help to eliminate high-energy barriers to substrate entry and product release that could have been caused by tighter packing of the TSA in higher activity HG variants.

### HG4, an efficient artificial enzyme

All of the structural features that enhance activity described above are caused primarily by residues within or close to the active site, which suggests that mutagenesis far from the active site may not be essential to create an efficient artificial enzyme. To test this hypothesis, we generated a variant of HG3 that contains all HG3.17 mutations found within 7.5 Å of the TSA, with the exception of N47E, which we omitted to favor the formation of a unit cell similar to that of HG3. We also included the R275A and W276F mutations found to widen the active site entrance. This yielded HG4, a variant of HG3 containing 8 mutations (Table 1, Supplementary Table 1). Kinetic analysis of HG4 revealed that its catalytic efficiency is >700-fold higher than that of HG3 (Table 1, Supplementary Figure 1), and equivalent to that of the average natural enzyme (~10^5^ M^−1^s^−1^).^22^ Crystallographic analysis of HG4 (Supplementary Tables 2–3) showed that its structure is highly similar to that of HG3.17 but with an active site that is better pre-organized (Figure 2–3, Supplementary Figure 3–4). Interestingly, both HG4 and HG3.17 have catalytic efficiencies on the order of 10^5^ M^−1^ s^−1^ despite the fact that the former enzyme contains less than half of the latter’s mutations, demonstrating that distal mutations in HG3.17 contribute little to its catalytic efficiency.

### Computational design of HG4

Given that all but one mutation (G82A) in HG4 are found at sites that were optimized during design of HG2,^5^ we investigated whether the HG4 structure could be accurately predicted using a computational protocol similar to the one that produced HG2 (Methods, Supplementary Tables 4–6). To do so, we first performed a positive control calculation in which rotamers for the HG4 sequence were optimized on the crystal structure backbone of TSA-bound HG4. This calculation yielded an *in silico* model of HG4 with an energy score and a predicted rotameric configuration in excellent agreement with the crystal structure (Figure 4a). This control demonstrates that the combination of energy function, rotamer library, and search algorithm used in this protocol is sufficiently accurate for recapitulating the structure of HG4, provided that the correct template, binding pose, and catalytic dyad is allowed. By contrast, when we replaced the HG4 backbone template with the *Thermoascus aurantiacus* xylanase 10A backbone used to design HG2 (PDB ID: 1GOR),^20^ we obtained a structural model that differs significantly from the HG4 crystal structure and that is destabilized by approximately 40 kcal/mol (Figure 4b). This result demonstrates that the 1GOR backbone template is not well-suited to accommodate the HG4 sequence, as evidenced by differences between the 1GOR-derived model and the HG4 crystal structure. Specifically, the backbone at position 83 is shifted by 1.1 Å in the HG4 crystal structure relative to its position in the 1GOR template, causing the transition state to adopt an alternate binding pose that minimizes steric clashes with Gly83, which is accompanied by repacking of several residues around the transition state, including Gln50. Use of our HG3 crystal structures with or without TSA as the design template causes similar, but less severe, structural and energetic effects (Figure 4c,d). However, when we optimized rotamers for the HG4 sequence on ensembles of backbone templates generated using molecular dynamics restrained by the HG3 diffraction data (Methods), we were able to recapitulate the correct transition-state binding mode on several individual ensemble members, with energies comparable to that of the HG4 crystal structure (Figure 4e–f, Supplementary Figure 5). These results highlight the impact of small backbone geometry variations on the computational predictions, and suggest that computational enzyme design with a crystallographically-derived backbone ensemble could obviate the need for directed evolution by allowing catalytically-competent sub-states to be sampled during the design procedure.

**Figure 4.**
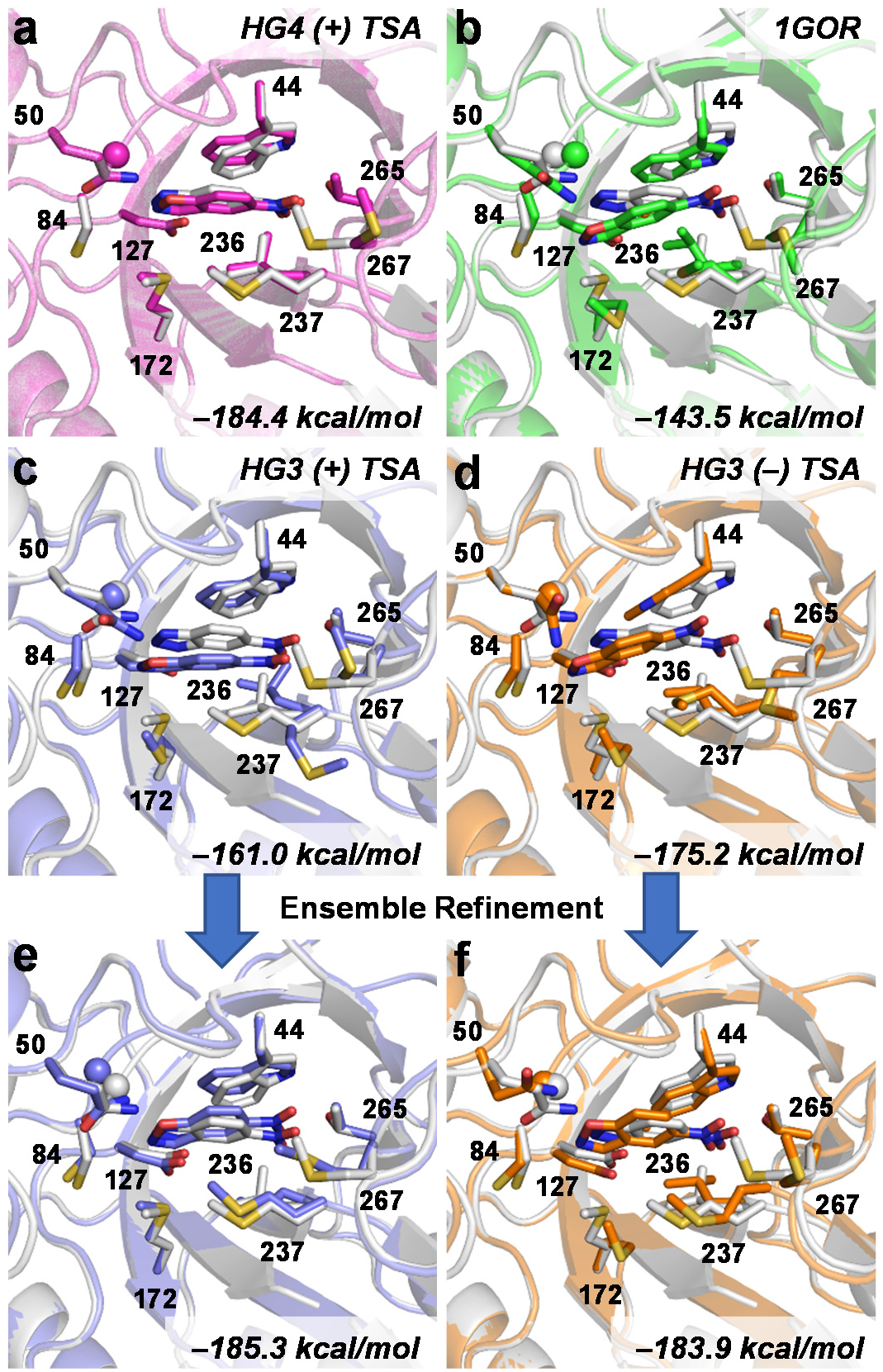
Computational design of HG4 on various backbone templates. The HG4 crystal structure with bound TSA (white) is overlaid on the HG4 design models (colored) obtained using the crystal structure of (a) HG4 with bound TSA, (b) *Thermoascus aurantiacus* xylanase 10A (PDB ID: 1GOR), (c) HG3 with bound TSA, or (d) HG3 without TSA. Panels (e) and (f) show the HG4 design models obtained using the template prepared by ensemble refinement from the corresponding HG3 density map that gave the best energy following repacking. PHOENIX energies of design models after repacking are indicated at bottom right. For reference, the energy of the HG4 crystal structure with bound transition state is –182.7 kcal/mol. In all cases, the transition state and transition state analogue are show at the center of barrel. Side chains of all residues forming the binding pocket are shown with the exception of Ala21 and Pro45, which were omitted for clarity. The sphere shows the alpha carbon of Gly83.

## Discussion

In this work, we followed changes to the conformational ensemble that occur during evolution of an enzyme with *de novo* biocatalytic function. Unlike previous examples where the active sites of *de novo* enzymes were completely remodeled during evolution,^23,24^ or where the binding pose of the substrate or transition state analogue was significantly altered,^13,17^ we observed only subtle changes to the active site geometry or TSA binding pose in the HG-series of Kemp eliminases. By contrast, many of the structural changes that contribute to enhanced catalysis in the HG series are dynamic in nature: the Gln50 catalytic residue became more rigid even though its average structure did not vary substantially, and the active site became better pre-organized via enrichment of catalytically-productive conformations of TSA-binding residues that were already present in the unbound state. These observations illustrate how small changes to the active site conformational ensemble can drive large changes in catalytic efficiency. Since these changes can be subtle and difficult to predict computationally, directed evolution can help increase activity by selecting for mutations that enrich catalytically-competent sub-states.^17,18^

Despite the challenges inherent to enzyme design, which are highlighted by our observations of the effects of mutations in the HG series of Kemp eliminases, our results suggest that *de novo* enzymes with native-like catalytic efficiencies can be computationally-designed, without the need to rely on subsequent improvement by laboratory directed evolution. Indeed, all mutations found in HG4 relative to the wild-type *Thermoascus aurantiacus* xylanase 10A template from which it is derived (PDB ID: 1GOR) are found at either first or second-shell residues, and these sites were all optimized during the original design of HG2.^5^ Yet, Privett *et al.* designed the lower activity enzyme HG2 instead of HG4. While Gln50 was not sampled as part of the catalytic dyad during design of HG2, the combination of the Asp127/Gln50 dyad with the productive transition-state binding pose would have scored poorly on the 1GOR template regardless. However, our approach to computational enzyme design that utilized an experimentally-derived ensemble of backbone templates yielded HG4 models with energies and binding modes comparable to that of the HG4 crystal structure. These results suggest an iterative approach to computational enzyme design that could circumvent the need for directed evolution by introducing an additional round of design that utilizes a backbone ensemble generated from experimental structural data obtained for an initial, low-activity enzyme. In the case of evolution, mutations are not selected for in the context of a single backbone conformation but instead across an entire conformational ensemble.^18^ Our ensemble design approach should therefore be more accurate than traditional approaches relying on a single backbone template because it allows the accessible conformational ensemble to be represented in the scoring of sequences. The incorporation of experimental restraints in the generation of the ensemble ensures that the computational procedure is applied to the true conformational ensemble that is sampled by the enzyme.

The results reported here provide additional support for the well-known fact that enzymes are plastic molecules whose backbone conformation can change upon introduction of mutations (as seen when comparing the 1GOR and HG-series crystal structures), and suggest improvements to the enzyme design protocol that can account for this property. This could be achieved by incorporating flexible backbone design algorithms during the repacking step,^25,26^ or by using pre-generated ensembles of energetically-accessible backbone templates,^27,28^ as was done here. While these methodological changes may improve the design of the enzyme transition state, it is likely that the creation of *de novo* enzymes with native-like catalytic efficiencies for more complex reactions will require a holistic approach where every possible state that the enzyme samples along its reaction coordinate is included in the design calculation. This could be achieved by the implementation of multistate approaches to computational protein design that allow the design of protein energy landscapes,^29^ rather than single structures. We expect that the structures reported here, especially those of HG4 and HG3, will be helpful to benchmark these future enzyme design protocols.

## Methods

### Protein expression and purification

Codon-optimized and his-tagged (C-terminus) genes for HG-series Kemp eliminases (Supplementary Table 1) cloned into the pET-11a vector (Novagen) via *Nde*I and *Bam*HI were obtained from Genscript. Enzymes were expressed in *E. coli* BL21-Gold (DE3) cells (Agilent) using lysogeny broth (LB) supplemented with 100 μg/mL ampicillin. Cultures were grown at 37 °C with shaking to an optical density at 600 nm of 0.3, at which point the incubation temperature was reduced to 18 °C. At an OD600 of 0.6, protein expression was initiated with 1 mM isopropyl β-D-1-thiogalactopyranoside. Following incubation for 16 hours at 18 °C with shaking (250 rpm), cells were harvested by centrifugation, resuspended in 10 mL lysis buffer (5 mM imidazole in 100 mM potassium phosphate buffer, pH 8.0), and lysed with an EmulsiFlex-B15 cell disruptor (Avestin). Proteins were purified by immobilized metal affinity chromatography according to the manufacturer’s protocol (Qiagen), followed by gel filtration in 50 mM sodium citrate buffer (pH 5.5) and 150 mM sodium chloride using an ENrich SEC 650 size-exclusion chromatography column (Bio-Rad). Purified samples were concentrated using Amicon Ultracel-10K centrifugal filter units (EMD Millipore).

### Steady-state kinetics

All assays were carried out at 27 °C in 100 mM sodium phosphate buffer (pH 7.0) supplemented with 100 mM sodium chloride. Triplicate reactions with varying concentrations of 5-nitrobenzisoxazole (AstaTech) dissolved in methanol (10% final concentration) were initiated by addition of approximately 2 μM HG3, 50 nM HG3.3b, 10 nM HG3.7/HG3.14, or 5 nM HG3.17/HG4. Product formation was monitored spectrophotometrically at 380 nm (ε = 15,800 M^−1^ cm^−1^).^5^ Linear phases of the kinetic traces were used to measure initial reaction rates. Initial reaction rates at different substrate concentrations were fit to the Michaelis-Menten equation using GraphPad Prism.

### Crystallization

Enzyme variants were prepared in 50 mM sodium citrate buffer (pH 5.5) at the concentrations listed in Supplementary Table 2. For samples that were co-crystallized with the transition state analog (TSA) 5-nitrobenzotriazole (AstaTech), a 100 mM stock solution of the TSA was prepared in dimethylsulfoxide (DMSO) and diluted 20-fold in the enzyme solutions for a final concentration of 5 mM TSA (5% DMSO). For each enzyme variant, we carried out initial crystallization trials in 15-well hanging drop format using EasyXtal crystallization plates (Qiagen) and a crystallization screen that was designed to explore the chemical space around the crystallization conditions reported by Blomberg *et al*.^15^ Crystallization drops were prepared by mixing 1 μL of protein solution with 1 μL of the mother liquor, and sealing the drop inside a reservoir containing an additional 500 μL of the mother liquor solution. The mother liquor solutions contained ammonium sulfate as a precipitant in sodium acetate buffer (100 mM), and the specific growth conditions that yielded the crystals used for X-ray data collection are provided in Supplementary Table 2. In some cases, a microseeding protocol was required to obtain high-quality crystals. Microseeds were prepared by vortexing crystals in their mother liquor in the presence of glass beads (0.5 mm), and were subsequently diluted into the mother liquor solutions used to form the crystallization drops.

### X-ray data collection and processing

Prior to X-ray data collection, crystals were mounted in polyimide loops and sealed using a MicroRT tubing kit (MiTeGen). Single-crystal X-ray diffraction data was collected on beamline 8.3.1 at the Advanced Light Source. The beamline was equipped with a Pilatus3 S 6M detector, and was operated at a photon energy of 11111 eV. Crystals were maintained at 277 K throughout the course of data collection. Each data set was collected using a total X-ray dose of 200 kGy or less, and covered a 180° wedge of reciprocal space. Multiple data sets were collected for each enzyme variant.

X-ray data was processed with the Xia2 program (https://doi.org/10.1107/S0021889809045701), which performed indexing, integration, and scaling with XDS and XSCALE,^30^ followed by merging with Pointless.^31^ For each variant, multiple individual data sets were merged to obtain the final set of reduced intensities, and the resolution cutoff was taken where the CC_1/2_ and <I/σI> values for the merged intensities fell to approximately 0.5 and 1.0 respectively. Information regarding data collection and processing is presented in Supplementary Table 3. The reduced diffraction data were analyzed with phenix.xtriage (http://www.ccp4.ac.uk/newsletters/newsletter43/articles/PHZ_RWGK_PDA.pdf) to check for crystal pathologies, and no complications were identified.

### Structure determination

We obtained initial phase information for calculation of electron density maps by molecular replacement using the program Phaser,^32^ as implemented in the PHENIX suite.^33^ Several different HG-series enzymes were used as molecular replacement search models. All members of the HG-series of enzymes crystallized in the same crystal form, containing two copies of the molecule in the crystallographic asymmetric unit, except for HG3.17, which crystallized with only one molecule in the asymmetric unit. To avoid model bias that could originate from using other members of the HG-series as molecular replacement search models, we applied random coordinate displacements (σ = 0.5 Å) to the atoms, and performed coordinate refinement against the structure factor data before proceeding to manual model building.

Next, we performed iterative steps of manual model rebuilding followed by refinement of atomic positions, atomic displacement parameters, and occupancies using a translation-libration-screw (TLS) model, a riding hydrogen model, and automatic weight optimization. All model building was performed using Coot^34^ and refinement steps were performed with phenix.refine (v1.13-2998) within the PHENIX suite.^33,35^ Restraints for the TSA were generated using phenix.elbow,^36^ starting from coordinates available in the Protein Data Bank (PDB ligand ID: 6NT).^37^ Further information regarding model building and refinement, as well as PDB accession codes for the final models, are presented in Supplementary Table 3. Time-averaged ensembles were generated for HG3 with and without ligand with phenix.ensemble_refinement implemented in PHENIX. To prepare the structures for ensemble refinement, low-occupancy conformers were removed, and occupancies adjusted to 100% using phenix.pdbtools. Hydrogen atoms were then added using phenix.ready_set. This procedure yielded 79- and 49-member ensembles from the HG3 structures with and without TSA, respectively.

### Computational enzyme design

All calculations were performed with the Triad protein design software (Protabit, Pasadena, CA, USA) using a Monte Carlo with simulated annealing search algorithm for rotamer optimization. The crystal structure of *Thermoascus aurantiacus* xylanase 10A was obtained from the Protein Data Bank (PDB code: 1GOR^20^) and further refined as described above to fix modeling issues with Thr84. Structures of HG3 with and without TSA, HG4 with TSA, and ensembles of HG3-derived templates were obtained from refinement of crystallographic data as described above. Following extraction of protein heavy-atom coordinates for the highest occupancy conformer from chain A, hydrogen atoms were added using the *addH.py* application in Triad. The Kemp elimination transition state (TS) structure^38^ was built using the parameters described by Privett and coworkers.^5^ Residue positions surrounding Asp127 were mutated to Gly (Supplementary Table 4), with the exception of position 50, which was mutated to Gln. A backbone-independent rotamer library^39^ with expansions of ± 1 standard deviation around χ1 and χ2 was used to provide side-chain conformations. A library of TS poses was generated in the active site by targeted ligand placement^2^ using the contact geometries listed in Supplementary Table 5. TS pose energies were calculated using the PHOENIX energy function,^5^ which consists of a Lennard-Jones 12–6 van der Waals term from the Dreiding II force field^40^ with atomic radii scaled by 0.9, a direction-dependent hydrogen bond term with a well depth of 8.0 kcal mol^−1^ and an equilibrium donor-acceptor distance of 2.8 Å,^41^ an electrostatic energy term modelled using Coulomb’s law with a distance-dependent dielectric of 10, an occlusion-based solvation potential with scale factors of 0.05 for nonpolar burial, 2.5 for nonpolar exposure, and 1.0 for polar burial,^42^ and a secondary structural propensity term.^43^ During the energy calculation step, TS–side-chain interaction energies were biased to favor interactions that satisfy contact geometries (Supplementary Table 6) as described by Lassila *et al*.^2^

Following ligand placement, the 10 lowest energy TS poses found on each template (HG4 with TSA, 1GOR, HG3 with TSA, and HG3 without TSA) were selected as starting points for repacking of the HG4 sequence. For individual members of the crystallographically-derived ensembles, only the single lowest energy TS pose was used for repacking. In the repacking calculation, the TS structure was translated ± 0.4 Å in each Cartesian coordinate in 0.2-Å steps, and rotated 10° about all three axes (origin at TS geometric center) in 5° steps for a total combinatorial rotation/translation search size of 5^6^ or 15,625 poses. Residues that were converted to Gly in the ligand placement step were allowed to sample all conformations of the amino acid found at that position in the HG4 sequence (Supplementary Table 4). The identities of the catalytic residues were fixed and allowed to sample all conformations of that amino-acid type. Side-chain– TS interaction energies were biased to favor those contacts that satisfy the geometries as done during the ligand placement step (Supplementary Table 6). Rotamer optimization was carried out using the search algorithm, rotamer library, and energy function described above. The single lowest energy repacked structure on each backbone template was used for analysis. To compare energies of the HG4 models obtained on the various templates, we calculated the energy difference between each repacked structure and the corresponding all-Gly structure obtained after ligand placement, and these energies are reported throughout the figures and text.

### Statistics and reproducibility

Experiments were repeated in triplicate where feasible. All replications were successful and the resulting data is presented with error values representing the standard deviation between replicates. No data was excluded from analyses.

## Data availability

Structure coordinates for all HG-series Kemp eliminases have been deposited in the Protein Data Bank with the following accession codes: HG3 (PDB ID: 5RG4, 5RGA), HG3.3b (PDB ID: 5RG5, 5RGB), HG3.7 (PDB ID: 5RG6, 5RGC), HG3.14 (PDB ID: 5RG7, 5RGD), HG3.17 (PDB ID: 5RG8, 5RGE), and HG4 (PDB ID: 5RG9, 5RGF).

## Acknowledgments

R.A.C. acknowledges an Early Researcher Award from the Ontario Ministry of Economic Development & Innovation (ER14-10-139), and grants from the Natural Sciences and Engineering Research Council of Canada (NSERC, RGPIN-2016-04831) and the Canada Foundation for Innovation (26503). J.S.F. is supported by a Searle Scholar Award from the Kinship Foundation, a Pew Scholar Award from the Pew Charitable Trusts, a Packard Fellowship from the David and Lucile Packard Foundation, NIH GM110580, UC Office of the President Laboratory Fees Research Program LFR-17-476732, and NSF STC-1231306. A.B. is the recipient of a postdoctoral fellowship from NSERC. M.C.T. is supported by a Ruth L. Kirschstein National Research Service Award (F32 HL129989). We thank Dr. Heidi K. Privett for providing HG2 design scripts and computational model, and Dr. Justin T. Biel for assistance with initial processing of X-ray datasets. Data collection at Beamline 8.3.1 at the Advanced Light Sources is supported by the University of California Office of the President, Multicampus Research Programs and Initiatives grant MR-15-328599, the Program Breakthrough Biomedical Research (which is partially funded by the Sandler Foundation), the National Institutes of Health (R01 GM124149 and P30 GM124169), Plexxikon Inc., and the Integrated Diffraction Analysis Technologies program of the US Department of Energy Office of Biological and Environmental Research. The Advanced Light Source (Berkeley, CA) is a national user facility operated by Lawrence Berkeley National Laboratory on behalf of the US Department of Energy under contract number DE-AC02-05CH11231, Office of Basic Energy Sciences.

## Author Contributions

A.B. and R.A.C. conceived the project. A.B., N.Z., E.N., N.M., and L.L. purified proteins. A.B. and N.Z. performed enzyme kinetics experiments. R.A.C. and M.C.T. crystallized proteins and performed X-ray diffraction experiments. A.B., R.V.R., M.C.T., and R.A.C. performed refinements. M.C.T. and J.S.F. designed X-ray crystallography experiments. A.B. and R.V.R. performed computational design experiments. R.A.C. wrote the manuscript. A.B., R.V.R., and M.C.T. edited the manuscript.

