## Supplementary Information for "Evolution of an enzyme conformational ensemble guides design of an efficient biocatalyst"

**This file includes:**

Supplementary Tables 1–6
Supplementary Figures 1–5

**Supplementary Table 1.** Amino-acid sequences of HG-series Kemp eliminases

| Enzyme | # mutations from HG3 | Sequence[a] |
|---|---|---|
| **HG3** | – | MAEAAQSVDQLIKARGKVYFGVATDQNRLTTGKNAAIIQADFGMVWPENS<br>MKWDATEPSQGNFNFAGADYLVNWAQQNGKLIGGGMLVWHSQLPSWVSSI<br>TDKNTLTNVMKNHITTLMTRYKGKIRAWDVVGEAFNEDGSLRQTVFLNVI<br>GEDYIPIAFQTARAADPNAKLYIMDYNLDSASYPKTQAIVNRVKQWRAAG<br>VPIDGIGSQTHLSAGQGAGVLQALPLLASAGTPEVSILMLDVAGASPTDY<br>VNVVNACLNVQSCVGITVFGVADPDSWRASTTPLLFDGNFNPKPAYNAIV<br>QDLQQGSIEGRGHHHHHH |
| **HG3.3b** | 6 | MAEAAQS**I**DQLIKARGKVYFGVATDQNRLTTGKNAAIIQADFGMVWPENS<br>M**H**WDATEPSQGNFNFAGADYLVNWAQQNGKLIGGG**C**LVWH**RD**LPSWVSSI<br>TDKNTLTNVMKNHITTLMTRYKGKIR**N**WDVVGEAFNEDGSLRQTVFLNVI<br>GEDYIPIAFQTARAADPNAKLYIMDYNLDSASYPKTQAIVNRVKQWRAAG<br>VPIDGIGSQTHLSAGQGAGVLQALPLLASAGTPEVSILMLDVAGASPTDY<br>VNVVNACLNVQSCVGITVFGVADPDSWRASTTPLLFDGNFNPKPAYNAIV<br>QDLQQGSIEGRGHHHHHH |
| **HG3.7** | 7 | MAEAAQS**I**DQLIKARGKVYFGVATDQNRLTTGKNAAII**K**ADFGMVWPENS<br>M**Q**WDATEPSQGNFNFAGADYLVNWAQQNGKLIGGG**C**LVWH**RH**LPSWVSSI<br>TDKNTLTNVMKNHITTLMTRYKGKIR**N**WDVVGEAFNEDGSLRQTVFLNVI<br>GEDYIPIAFQTARAADPNAKLYIMDYNLDSASYPKTQAIVNRVKQWRAAG<br>VPIDGIGSQTHLSAGQGAGVLQALPLLASAGTPEVSILMLDVAGASPTDY<br>VNVVNACLNVQSCVGITVFGVADPDSWRASTTPLLFDGNFNPKPAYNAIV<br>QDLQQGSIEGRGHHHHHH |
| **HG3.14** | 12 | MAEAAQS**I**DQLIKARGKVYFGVATDQNRLTTGKNAAII**K**ADFGMVWPENS<br>M**Q**WDATEPSQGNFNFAGADYLVNWAQQNGKLIG**A**G**C**LVWHS**H**LPSWVSSI<br>TDKNTL**I**NVMKNHITTLMTRYKGKIR**T**WDVVGEAFNEDGSLRQ**N**VFLNVI<br>GEDYIPIAFQTARAADPNAKLYIMDYNLDSASYPKTQAIVNRVKQWRAAG<br>VPIDGIGSQ**M**HLSAGQGAGVLQALPLLASAGTPEVSILMLDVAGASPTDY<br>VNVVNACLNVQSCVGITVFGVADPDSWRAS**S**TPLLFDGNFNPKPAYNAIV<br>Q**N**LQQGSIEGRGHHHHHH |
| **HG3.17** | 17 | MAEAAQS**I**DQLIKARGKVYFGVATDQNRLTTGKNAAII**K**ADFGMVWPE**E**S<br>M**Q**WDATEPSQGNFNFAGADYLVNWAQQNGKLIG**A**G**C**LVWH**NF**LPSWVSSI<br>TDKNTL**I**NVMKNHITTLMTRYKGKIR**T**WDVVGEAFNEDGSLRQ**N**VFLNVI<br>GEDYIPIAFQTARAADPNAKLYIMDYNLDSASYPKTQAIVNRVKQWRAAG<br>VPIDGIGSQ**M**HLSAGQGAGVLQALPLLASAGTPEVSILMLDVAGASPTDY<br>VNVVNACLNVQSCVGITV**M**GVADPDS**AF**AS**S**TPLLFDGNFNPKPAYNAIV<br>Q**N**LQQGSIEGRGHHHHHH |
| **HG4** | 8 | MAEAAQSVDQLIKARGKVYFGVATDQNRLTTGKNAAIIQADFGMVWPENS<br>M**Q**WDATEPSQGNFNFAGADYLVNWAQQNGKLIG**A**G**C**LVWHS**F**LPSWVSSI<br>TDKNTLTNVMKNHITTLMTRYKGKIR**T**WDVVGEAFNEDGSLRQTVFLNVI<br>GEDYIPIAFQTARAADPNAKLYIMDYNLDSASYPKTQAIVNRVKQWRAAG<br>VPIDGIGSQTHLSAGQGAGVLQALPLLASAGTPEVSILMLDVAGASPTDY<br>VNVVNACLNVQSCVGITV**M**GVADPDS**AF**ASTTPLLFDGNFNPKPAYNAIV<br>QDLQQGSIEGRGHHHHHH |

[a] Mutations from HG3 are highlighted in bold. All sequences contain a His-tag at the C-terminus.

**Supplementary Table 2.** Crystallization conditions

| Enzyme [a] | TSA [b] | pH | Protein (mg mL$^{-1}$) | $(NH_4)_2SO_4$ (M) |
|---|---|---|---|---|
| **HG3** | (–) | 4.6 | 4 | 1.8 |
| | (+) | 4.6 | 4 | 1.8 |
| **HG3.3b** | (–) | 5.4 | 5 | 2.0 |
| | (+) | 5.4 | 5 | 2.0 |
| **HG3.7** | (–) | 4.0 | 6 | 1.6 |
| | (+) | 5.0 | 12 | 2.0 |
| **HG3.14** | (–) | 4.5 | 3 | 1.6 |
| | (+) | 5.0 | 3 | 2.4 |
| **HG3.17** | (–) | 5.0 | 5 | 1.6 |
| | (+) | 5.0 | 10 | 1.6 |
| **HG4** | (–) | 4.6 | 10 | 1.6 |
| | (+) | 4.6 | 10 | 1.6 |

[a] All enzymes were crystallized in 100 mM sodium acetate buffer at the indicated pH.
[b] TSA (5-nitrobenzotriazole) was dissolved in pure DMSO and added to crystallization solution at a final concentration of 5 mM (5% DMSO).

**Supplementary Table 3.** Crystallographic data and refinement statistics for room-temperature structures (277 K)

| | **HG3** | | **HG3.3b** | | **HG3.7** | | **HG3.14** | | **HG3.17** | | **HG4** | |
|---|---|---|---|---|---|---|---|---|---|---|---|---|
| **TSA** | (–) | (+) | (–) | (+) | (–) | (+) | (–) | (+) | (–) | (+) | (–) | (+) |
| PDB ID | 5RG4 | 5RGA | 5RG5 | 5RGB | 5RG6 | 5RGC | 5RG7 | 5RGD | 5RG8 | 5RGE | 5RG9 | 5RGF |
| **Data collection** [a] | | | | | | | | | | | | |
| Resolution (Å) | 41.52–1.99 | 55.15–1.89 | 37.09–1.62 | 48.16–1.44 | 60.34–1.35 | 79.81–1.39 | 79.83–1.47 | 48.94–1.40 | 46.17–1.73 | 38.46–1.77 | 79.99–1.47 | 41.50–1.40 |
| Space group | $P2_12_12_1$ | $P2_12_12_1$ | $P2_12_12_1$ | $P2_12_12_1$ | $P2_12_12_1$ | $P2_12_12_1$ | $P2_12_12_1$ | $P2_12_12_1$ | $P2_12_12_1$ | $P2_12_12_1$ | $P2_12_12_1$ | $P2_12_12_1$ |
| *Cell params.* | | | | | | | | | | | | |
| a b c (Å) | 76.14 79.97 99.06 | 76.24 79.88 99.05 | 76.23 80.03 98.91 | 76.24 79.92 98.71 | 76.20 79.85 98.81 | 76.26 79.81 98.72 | 76.31 79.83 99.11 | 76.32 79.81 98.94 | 51.35 58.17 92.35 | 50.90 57.98 95.69 | 76.29 79.99 98.98 | 75.97 77.99 98.03 |
| α β γ (°) | 90 90 90 | 90 90 90 | 90 90 90 | 90 90 90 | 90 90 90 | 90 90 90 | 90 90 90 | 90 90 90 | 90 90 90 | 90 90 90 | 90 90 90 | 90 90 90 |
| Molecules per asymm. unit | 2 | 2 | 2 | 2 | 2 | 2 | 2 | 2 | 1 | 1 | 2 | 2 |
| $R_{pim}$ | 0.125 (0.375) | 0.067 (0.332) | 0.047 (0.407) | 0.043 (0.540) | 0.029 (0.246) | 0.029 (0.359) | 0.029 (0.372) | 0.046 (0.391) | 0.088 (0.734) | 0.101 (0.606) | 0.048 (0.550) | 0.028 (0.375) |
| $CC_{1/2}$ | 0.974 (0.679) | 0.994 (0.745) | 0.997 (0.690) | 0.998 (0.524) | 0.998 (0.826) | 0.999 (0.681) | 0.999 (0.586) | 0.998 (0.697) | 0.996 (0.587) | 0.990 (0.499) | 0.997 (0.609) | 0.999 (0.743) |
| I/σI | 3.7 (1.0) | 6.8 (1.0) | 8.9 (1.0) | 8.9 (1.1) | 13.0 (2.3) | 12.6 (1.3) | 13.6 (1.4) | 9.4 (1.4) | 8.8 (1.1) | 5.5 (1.0) | 8.2 (1.1) | 29.4 (3.5) |
| Completeness (%) | 100.0 (100.0) | 100.0 (100.0) | 100.0 (100.0) | 99.8 (99.8) | 99.0 (91.8) | 100.0 (99.6) | 99.8 (99.2) | 100.0 (100.0) | 100.0 (99.5) | 99.5 (99.6) | 99.7 (99.6) | 99.8 (99.0) |
| Multiplicity | 6.4 (6.5) | 12.7 (9.3) | 12.9 (10.9) | 6.4 (6.4) | 6.3 (4.7) | 12.7 (10.0) | 18.5 (10.4) | 13.0 (13.5) | 16.5 (6.3) | 6.5 (6.1) | 5.8 (5.9) | 17.3 (13.0) |
| # reflections (total unique) | 42219 | 51523 | 118990 | 113916 | 131151 | 121461 | 103204 | 80367 | 29625 | 27504 | 103164 | 114707 |
| **Refinement** | | | | | | | | | | | | |
| R work/free | 0.1767/ 0.2164 | 0.1482/ 0.1863 | 0.1421/ 0.1665 | 0.1360/ 0.1665 | 0.1271/ 0.1412 | 0.1255/ 0.1409 | 0.1319/ 0.1480 | 0.1329/ 0.1518 | 0.1397/ 0.1615 | 0.1501/ 0.1794 | 0.1434/ 0.1633 | 0.1317/ 0.1497 |
| *No. atoms* | | | | | | | | | | | | |
| Protein | 4754 | 5017 | 5261 | 5376 | 5661 | 5703 | 5407 | 5066 | 2398 | 2387 | 5164 | 5180 |
| Ligand | – | 24 | – | 24 | – | 24 | – | 24 | – | 12 | – | 24 |
| Water | 424 | 483 | 463 | 498 | 543 | 527 | 511 | 496 | 267 | 228 | 474 | 488 |
| *RMSD* | | | | | | | | | | | | |
| bond lengths (Å) | 0.005 | 0.008 | 0.004 | 0.005 | 0.008 | 0.008 | 0.008 | 0.005 | 0.004 | 0.003 | 0.004 | 0.004 |
| bond angles (°) | 0.831 | 0.910 | 0.784 | 0.880 | 1.029 | 0.986 | 0.973 | 0.835 | 0.731 | 0.631 | 0.773 | 0.822 |
| MolProbity clashscore | 2.97 | 3.09 | 3.35 | 4.67 | 3.89 | 2.02 | 2.32 | 2.26 | 1.26 | 1.26 | 2.91 | 2.70 |

[a] Highest resolution shell is shown in parentheses.

**Supplementary Table 4.** Amino-acid positions optimized during computational design of HG4

| Ligand placement [a] | Repacking [b] |
|---|---|
| 16, 17, 21, 42, 44, 46, 47, 79, 81, 83, 84, 87, 90, 125, 130, 170, 172, 207, 209, 234, 236, 237, 239, 267, 275, 276 | V16, Y17, A21, M42, W44, E46, N47, Q50, L79, G81, A82, G83, C84, W87, F90, T125, D127, G130, Y170, M172, Q207, H209, S234, L236, M237, D239, M267, A275, F276 |

[a] Positions that were mutated to Gly during ligand placement. Catalytic residues D127 and Q50 were allowed to sample alternate rotamers. All other residues were kept fixed.
[b] Positions and amino-acid types that were allowed to sample alternate rotamers during repacking. All other residues were kept fixed.

**Supplementary Table 5.** Geometric definitions for generation of transition-state poses off the side chains of catalytic residues

| Contact | Type | Atom 1 [a] | Atom 2 [a] | Atom 3 [a] | Atom 4 [a] | Values [b] |
|---|---|---|---|---|---|---|
| **Asp127** | Distance | OD1 or OD2 | **H3** | | | 1.0, 1.2, 1.5 |
| | Angle | CG | OD1 or OD2 | **H3** | | 112, 117, 122 |
| | Angle | OD1 or OD2 | **H3** | **C3** | | 159, 164, 169, 174, 179 |
| | Torsion | CB | CG | OD1 or OD2 | **H3** | 0, 5, 10, 170, 175, 180 |
| | Torsion | CG | OD1 or OD2 | H3 | **C3** | 170, 175, 180, 185, 190 |
| | Torsion | OD1 or OD2 | **H3** | **C3** | **N2** | 0, 5, 170, 175, 180 |
| **Gln50** | Distance | 1HE2 or 2HE2 | **O1** | | | 1.2, 1.5, 1.7, 1.9, 2.1, 2.3 |
| | Angle | NE2 | 1HE2 or 2HE2 | **O1** | | 145, 148, 151, 154, 157 |
| | Angle | 2HE2 | **O1** | **N2** | | 120, 125, 135, 145, 155 |
| | Torsion | CD | NE2 | 1HE2 or 2HE2 | **O1** | 115, 120, 135, 140, 145 |
| | Torsion | NE2 | 1HE2 or 2HE2 | **O1** | **N2** | 180, 190, 200, 210, 220, 230 |
| | Torsion | 1HE2 or 2HE2 | **O1** | **N2** | **C3** | 150, 160, 170, 180, 190, 200 |

[a] Atoms in bold are from the transition state. All other atoms are from the catalytic residues.
[b] Distance measurements given in Å, all others in degrees.

**Supplementary Table 6.** Geometric constraints used to define catalytic contacts during repacking step of HG4 computational design

| Contact | Type | Atom 1[a] | Atom 2[a] | Atom 3[a] | Atom 4[a] | Min[b] | Max[b] |
|---|---|---|---|---|---|---|---|
| **Asp127** | Distance | OD1 or OD2 | **H3** | | | 1.0 (1.0) | 1.6 (1.6) |
| | Angle | CG | OD1 or OD2 | **H3** | | 110 (110) | 130 (130) |
| | Angle | OD1 or OD2 | **H3** | **C3** | | 160 (160) | 179 (179) |
| | Torsion | CB | CG | OD1 or OD2 | **H3** | –20 (–20) | 20 (20) |
| **Gln50** | Distance | 1HE2 or 2HE2 | **O1** | | | 1.2 (1.2) | 2.3 (3.2) |
| | Angle | NE2 | 1HE2 or 2HE2 | **O1** | | 145 (131) | 157 (179) |
| | Angle | 1HE2 or 2HE2 | **O1** | **N2** | | 119 (112) | 139 (150) |
| | Torsion | 1HE2 or 2HE2 | **O1** | **N2** | **C3** | 161 (129) | 199 (199) |

[a] Atoms in bold are from the transition state. All other atoms are from the catalytic residues.
[b] Distance measurements given in Å, all others in degrees. Values in parentheses are for the repacking step, while the others are for ligand placement.

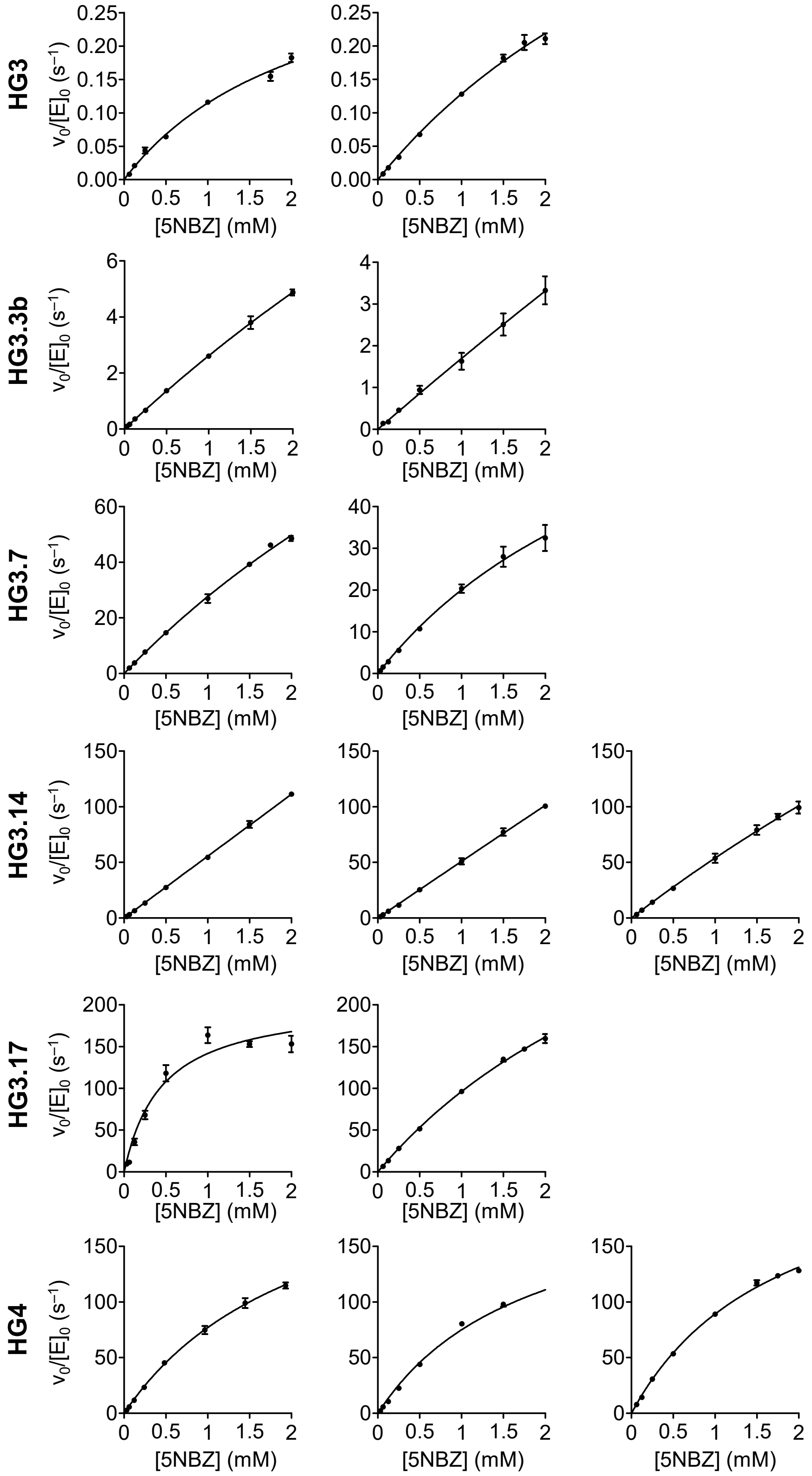

**Supplementary Figure 1. Steady-state kinetics.** Michaelis–Menten plots of normalized initial rates as a function of 5-nitrobenzisoxazole (5NBZ) concentrations are shown. All experiments were performed in triplicate (mean ± s.d.) on at least two independent enzyme samples.

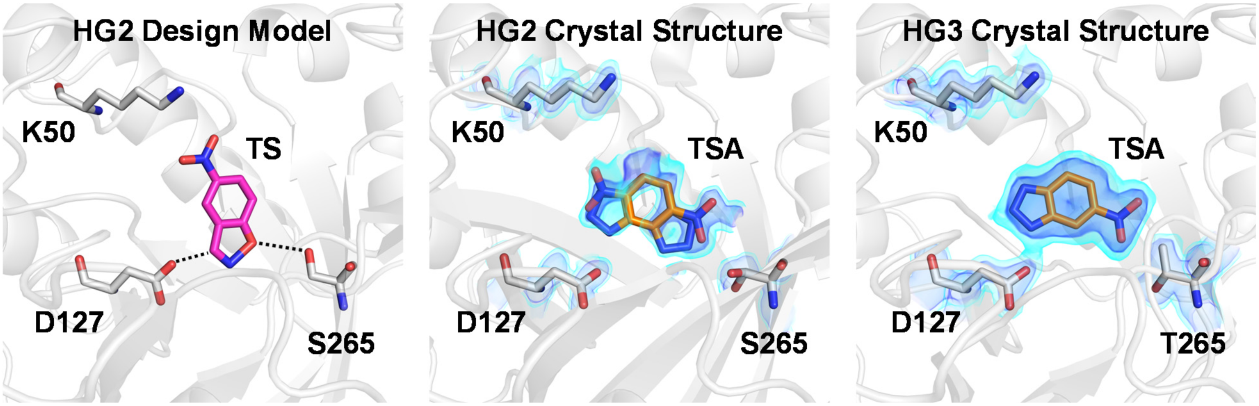

**Supplementary Figure 2. HG2 is the direct precursor to HG3.** For the crystal structures of HG2 and HG3, the 2Fo-Fc map is shown in volume representation at two contour levels: 0.5 eÅ$^{-3}$ and 1.5 eÅ$^{-3}$ in light and dark blue, respectively. HG2 was originally designed to stabilize the transition state (TS) via catalytic contacts (dashed lines) with the D127 base and the S265 hydrogen bond donor (Left panel). However, its crystal structure (PDB ID: 3NYD) showed that the transition state analogue (TSA) was bound in two alternate orientations (Middle panel). In the catalytically productive pose, the acidic N–H bond of the TSA that mimics the cleavable C–H bond of the substrate is located within H-bonding distance to the carboxylate oxygen of D127, and the nitro group is close to S265. In the catalytically non-productive pose, the TSA is flipped, which positions its nitro group closer to K50, and its acidic N–H bond far from the side chain of D127. To increase activity, Privett *et al.* introduced the S265T mutation into HG2, leading to HG3 (Right panel). This mutation was predicted by molecular dynamics to rigidify the active site, and thereby increase activity. Of note, no density for the non-productive binding pose of the TSA was observed in the HG3 crystal structure reported here.

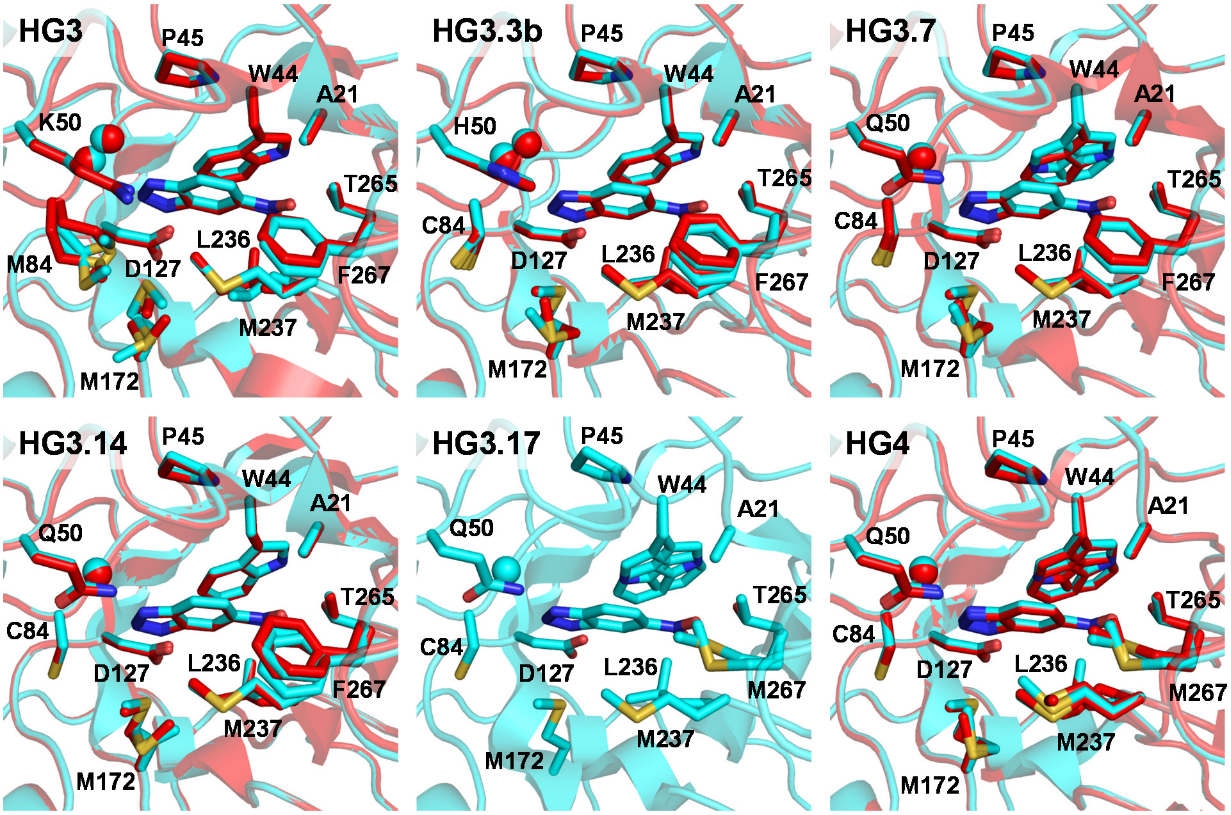

**Supplementary Figure 3. TSA-binding pocket.** Overlay of chains A (cyan) and B (red) showing residues forming the TSA-binding pocket in all HG-series Kemp eliminases. For HG3.17, only chain A is shown as its asymmetric unit contained a single polypeptide chain. In all cases, the TSA is bound in the middle of the barrel. Spheres indicate alpha carbons of Gly83 (two conformers, *cis*/*trans*, are observed for peptide bond between Gly83 and Met/Cys84 in HG3 and HG3.3b).

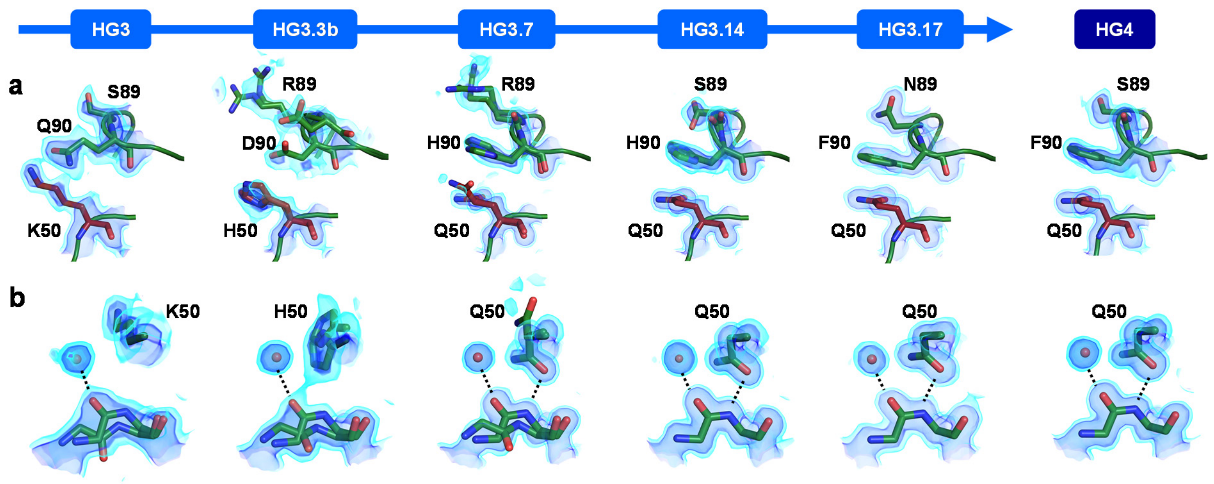

**Supplementary Figure 4. Crystal structures of HG-series Kemp eliminases in the unbound state.** In all cases, only atoms from chain A are shown. The 2Fo-Fc maps are shown in volume representation at two contour levels: 0.5 eÅ$^{-3}$ and 1.5 eÅ$^{-3}$ in light and dark blue, respectively. (a) Conformational changes to loop formed by residues 87–90 over the course of the evolutionary trajectory. (b) The peptide bond between residues 83 and 84 adopts both *cis* and *trans* conformations in HG3, HG3.3b, and HG3.7, but only the *cis* conformation in the higher activity variants. Ordered water molecules are shown as red spheres, and hydrogen bonds as dashed lines.

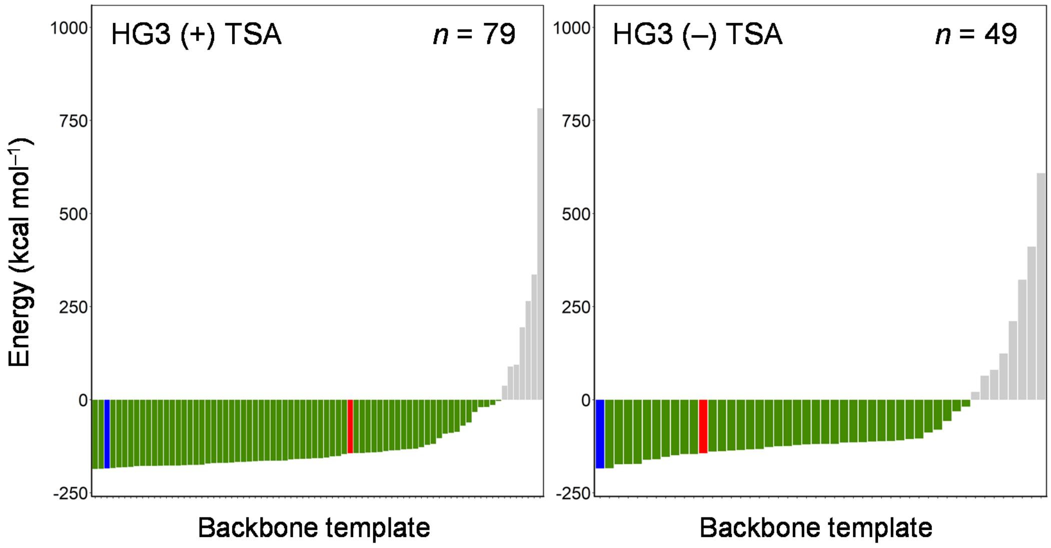

**Supplementary Figure 5. Energy of HG4 design models generated on various backbone templates.** Rotamers for the HG4 sequence and its associated transition state binding pose were optimized (Methods) on individual backbone templates (bars). These ensembles of backbone templates were generated using molecular dynamics fitted to the HG3 diffraction data with and without TSA. In all cases, *n* indicates the number of templates per ensemble. Green and grey bars indicate templates that yielded design models with favorable or unfavorable energy, respectively. Blue and red bars indicate design models obtained from the HG4 with bound TSA (–184.4 kcal/mol) or 1GOR (–143.5 kcal/mol) crystal structures, respectively. Several templates from each ensemble yielded HG4 design models with more favorable energy than that obtained on the 1GOR template, but only the HG3 with TSA ensemble yielded models that scored more favorably than the one obtained on the HG4 (+) TSA crystal structure.
